# Water storage and irrigation practices associated with cannabis production drive seasonal patterns of water extraction and use in Northern California watersheds

**DOI:** 10.1101/618934

**Authors:** Christopher Dillis, Connor McIntee, Ted Grantham, Van Butsic, Lance Le, Kason Grady

**Affiliations:** California State Water Resources Control Board, North Coast Region, Santa Rosa, California, United States of America; University of California Berkeley, Berkeley, California, United States of America

## Abstract

Concerns have been raised over the impacts of cannabis farms on the environment and water resources in particular, yet data on cultivation practices and water use patterns and have been limited. Estimates of water use for cannabis cultivation have previously relied on extrapolated values of plant water demand, which are unable to account for differences in cultivation practices, variation across the growing season, or the role of water storage in altering seasonal extraction patterns. The current study uses data reported by enrollees in California’s North Coast Regional Water Quality Control Board (Regional Water Board) Cannabis Program to model how variation in cultivation practices and the use of stored water affect the timing and amount of water extracted from the environment. We found that the supplemental use of stored water resulted in a seasonal pattern of water extraction (i.e. water withdrawals from the environment) that was distinct from water demand (i.e. water applied to plants). Although water input to storage in the off-season months (November through March) reduced water extraction in the growing season (April through October), farms generally did not have sufficient storage to completely forbear from surface water extraction during the growing season. Beginning in 2019, forbearance will be required during this period for those in the regulated cannabis industry. The two most important predictors of storage sufficiency (type of storage infrastructure and seasonality of water source) also had reliable effects on seasonal extraction patterns, further emphasizing the link between water storage and extraction profiles. These findings suggest that resource managers and policy makers should consider the ways in which cultivation practices drive water extraction patterns and how these practices may be influenced by participation in the regulated cannabis industry.

## Introduction

Northern California has long been the center of cannabis production in the United States [1–3]. Cannabis cultivation sites are distributed throughout the region and are generally located in remote, upper watersheds [4]. When the state voted to permit recreational cannabis use in 2016, a key argument for legalization was allowing the state to better address environmental harms caused by cannabis production [5]. In California, Illegal cannabis farms have been shown to fragment forested landscapes [6], introduce pesticides, fertilizers, and rodenticides into the environment [5, 7-9], and are often located in sensitive habitats, including along streams that support endangered salmon species [4].

There has been particular concern over the impacts of cannabis cultivation on water resources in areas with seasonally dry conditions [5]. Because many cannabis farms in Northern California are located in rural landscapes with no access to municipal water supplies, cultivators generally obtain water directly from the environment, relying on local springs, streams, and groundwater wells [10]. Stream flow has been identified as an important limiting factor to salmon, and other sensitive aquatic species in the region, particularly given the seasonal drought of California’s Mediterranean Climate [11–14]. Because cannabis water demands coincide with the summer dry season, agricultural water diversions in the North Coast Region have the potential to reduce stream flows [15–16], increase stream temperatures [17], or even dewater streams during critical life stages of aquatic species [18–19]. Although these streams are highly sensitive to variability in flow rates [20–21], there is a dearth of information surrounding cannabis water use practices, making it difficult to quantify potential environmental impacts.

An accurate baseline assessment of water use by cannabis cultivation is particularly important when considering the spatial and temporal distribution of cannabis water demands. Although cannabis cultivation has a relatively small geographic footprint, there is a high degree of spatial clustering among cultivation sites [6] at both local [22] and regional scales [4]. Currently, there are very few data on the cumulative impacts of many, dispersed water users [23–24] or flow estimates for small, unnamed streams on which they occur [25]. Impacts from densely clustered cannabis farms may be exacerbated by temporal clustering of water demand, with cannabis plants requiring frequent watering in late summer drought months and thus causing concern for instream flows [5, 18]. A key assumption behind this concern has been that water demand of cannabis plants directly results in water extraction during this period; however, there has been no systematic analysis of when water is drawn from the watershed or the factors that contribute to extraction patterns.

To date, estimates of water use by cannabis cultivation have relied on scaling a static approximation of outdoor cannabis plant demands during the growing season for outdoor cultivation, June-October [26–27, 18]. Unfortunately, this approach cannot account for changing water demands over the course of the growing season or under different cultivation conditions. For instance, a substantial proportion of farms use mixed-light operations (whether in greenhouses or “hoophouses”) that alter light cycles to produce multiple harvests of smaller cannabis plants, potentially extending the growing season, yet resulting in much lower water demand per-plant relative to outdoor cultivation. Another significant shortcoming of plant-based estimates is that they do not account for the practice of using stored water. Although cannabis farms are known to often utilize water storage, to date, detailed data on capacities have been sparse, given limited site access and the difficulty of obtaining these data from aerial imagery [28–29]. An improved estimation of the water demand of cannabis cultivation would account for water that is extracted and stored outside of the growing season, as well as how factors such as water sources shape both when and how much water is extracted and stored. These seasonal patterns of water extraction hold tremendous importance, given the potential for overlap between cannabis water demands and low summer water availability.

This study analyzed self-reported data from cannabis farmers that were enrolled for regulatory coverage under California’s North Coast Regional Water Quality Control Board Cannabis Waste Discharge Regulatory Program [10]. The reports were filtered to reduce bias and then analyzed through the development of multiple models that related water use practices to farm characteristics, including cultivation area, the type of operation (i.e. outdoor vs. mixed-light), water storage capacity, type of storage, and water source, to address the following questions:

1. Are water extraction rates distinct from those of water use (i.e. based on plant demand) over the growing season and are these patterns influenced by operation types?
2. Do farms typically have sufficient capacity to maintain a positive water storage balance for the entirety of the dry season (April through October) and what are the most important predictors of sufficiency?
3. How do the factors that influence water storage in turn affect the timing and amount of water extraction?

## Methods

### Data

The data used in this study were collected from cannabis farms enrolled for regulatory coverage under the North Coast Regional Water Quality Control Board Cannabis Waste Discharge Regulatory Program (NCRWQCB Cannabis Program). This program was established in August 2015, with the majority of enrollees entering the program in late 2016 and early 2017. The data discussed herein were collected from annual reports submitted in 2018 (n=1,702) and were required to reflect site conditions during the 2017 cultivation year. These data, therefore, largely represent the first full season of cultivation regulated by the NCRWQCB for the majority of enrollees in the Cannabis Program. Parcels with cannabis cultivation (including multiple contiguous parcels under a single ownership) constituted a *farm*, and reporting was done at this scale. Although the spatial extent of the NCRWQCB Cannabis Program included all of California’s North Coast Region, due to constraints placed on cultivation by local and county ordinances, reports from enrolled farms were limited to Humboldt, Trinity, Mendocino, and Sonoma Counties (Fig 1).

**Fig 1.**
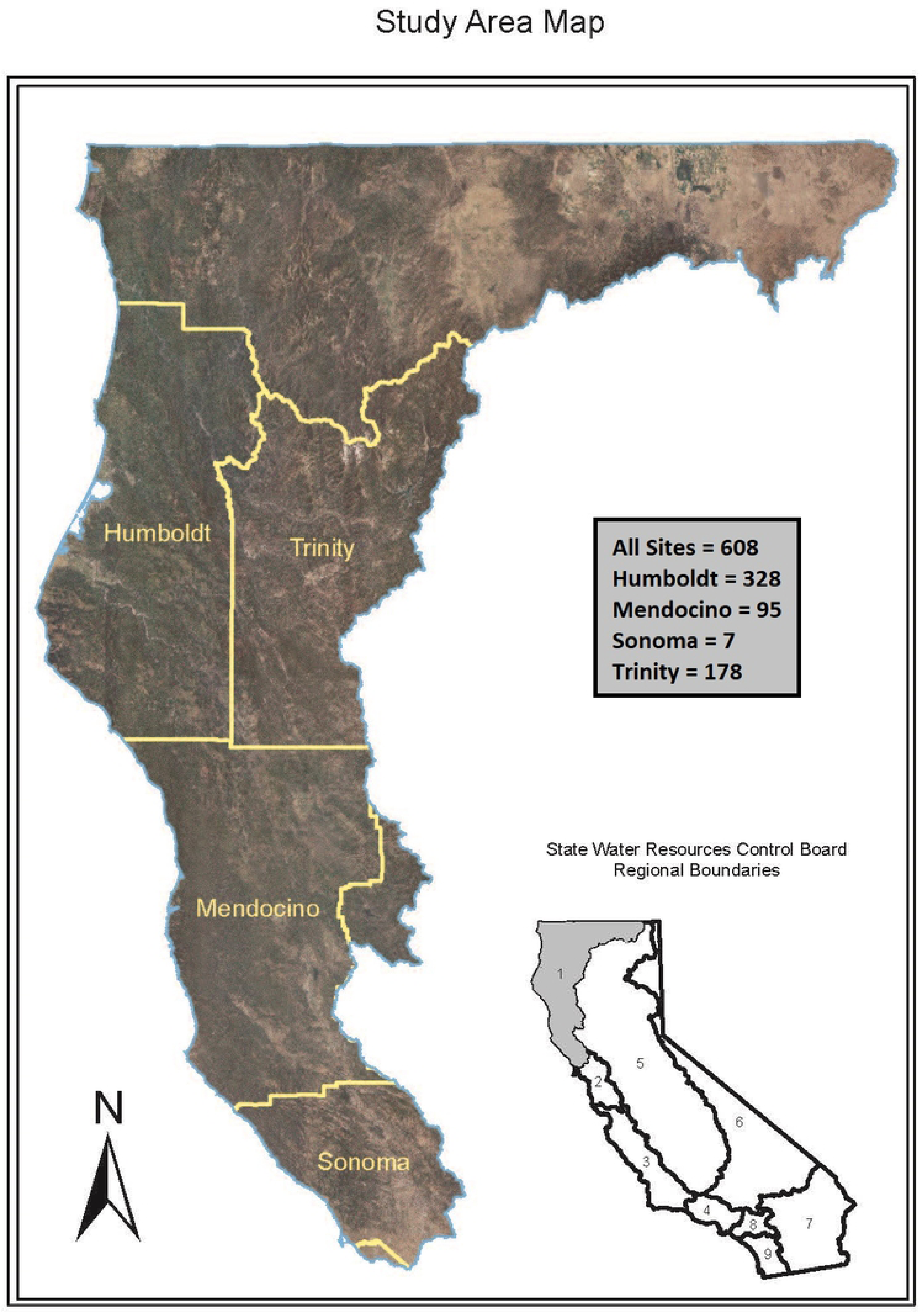
Study Area Map. The North Coast Region of California contains additional counties besides Humboldt, Trinity, Mendocino, and Sonoma; however, enrollments in the NCRWQCB Cannabis Program, and thus data included in the current study, were limited to these counties.

Given that data were self-reported, we screened reports for quality and excluded those that were not prepared by professional consultants. Additional criteria for excluding reports included: reported water applied from storage without any corresponding input to storage, substantial water input reported from “rain” during summer drought months, and failure to list a proper water source. Farms were not required to use water meters, and those without meters often made estimates based the frequency of filling and emptying of small temporary storage tanks (250 – 2500 gallons; 946 – 9,460 L) used for gravity feed systems and/or nutrient mixing. We attempted to identify and exclude farms with erroneous reporting by removing extreme water extraction outliers (more than 1.5 x Interquartile Range) and those with imprecise monthly estimates (e.g., 20,000, 25,000, and 30,000 L). Farms with total cultivation area over one acre (43,560 ft^2^; 4,046 m^2^) were also excluded, to minimize additional error inflation resulting from water use estimates at large (and infrequently occurring) scales. Farms that reported no water use for the entire season or no cultivation area were excluded from the analysis (reports were required from all enrollees regardless of whether cultivation occurred during 2017 season). Farms that reported a cultivation area of exactly 10,000 ft^2^ or 9,999 ft^2^ (929 or 928 m^2^, respectively) were determined to reflect regulatory thresholds for local cultivation ordinances rather than true cultivation area size. Aerial imagery from the National Agriculture Imagery Program (2016 NAIP) was reviewed to provide an improved estimate of the size of cultivation area for these farms. The final dataset included 608 reports.

The data reported for each farm included the size of cultivation area (*cultivation area*: ft^2^), volume of water applied to plants (*water applied*: gallons), volume of water input to storage (*water input*: gallons), type and volume of water storage infrastructure (*storage type*: pond, other (i.e. tank or water bladder); *storage capacity*: gallons). Although the data were reported on the standard measurement system, for the purposes of data analysis and reporting, these measures were converted to SI units. Water data were reported on a monthly basis (*month*), specifying up to three sources of applied water (*application source*: delivery, municipal, pond, rain, springs, surface, tanks, water bladder, or well) and water input to storage (*input source*: delivery, municipal, rain, springs, surface, or well). An additional parameter (*source type*: seasonal, perennial) was created specifying whether farms relied exclusively on seasonal water sources (e.g. rain, springs, surface) or had at least one perennial source (i.e. incorporating well, delivery, or municipal water). Although farms may have perennial access to springs and surface water, these water sources are subject to pending regulatory restrictions, which will prohibit water diversions from April through October (i.e. “forbearance period”). However, for the 2017 cultivation year, farms that reported use of these sources during this period were not subject to regulatory violations or penalties, nor did the use of these sources make them ineligible for enrollment [10].

Cultivation area was reported as the footprint of mixed-light infrastructure and outdoor gardens, incorporating both canopy area and the space between plants. Aerial imagery (2016 NAIP) was used to distinguish which farms had outdoor gardens, mixed-light infrastructure, or both, and an additional model parameter was created (*operation type*: outdoor, mixed-light, combination). The purpose of identifying *operation type* was to control for variation in plant spacing (plants under mixed-light cultivation are smaller and more tightly spaced), ambient temperature and humidity (mixed-light cultivation occurs underneath a canopy covering), and length of cultivation season (mixed-light operations tend to produce multiple harvests, although on shorter intervals).

### Seasonality of Water Use and Water Extraction

Water use and water extraction totals for each month were created using combinations of reported water applied to plants and water input to storage and these served as response variables for model fitting. W*ater use* was defined as water applied either from storage or directly applied from the original source, thus reflecting plant demand. *Water extraction* was defined as water either input to storage or directly applied from the original source, thus reflecting withdrawal from the watershed.

As an additional check of these self-reported data, we sought to compare the reported rates of *water use* against the commonly adopted figure of 22 L/plant/day [26–27, 18] for outdoor plants during the growing season. This was done using aerial imagery data from an existing study in which the number of mature cannabis plants were counted within a cultivation area of a known size to determine the size of cultivation area representative of a single, outdoor, cannabis plant. This number was determined to be 15 m^2^ of cultivation area, accounting for both the canopy of the plant and spacing between individuals. The rate of 22 L/ 15 m^2^/day was converted to 22 L/ 15 m^2^/month and depicted where appropriate (Figs 2 and 4) to provide context.

**Fig 2.**
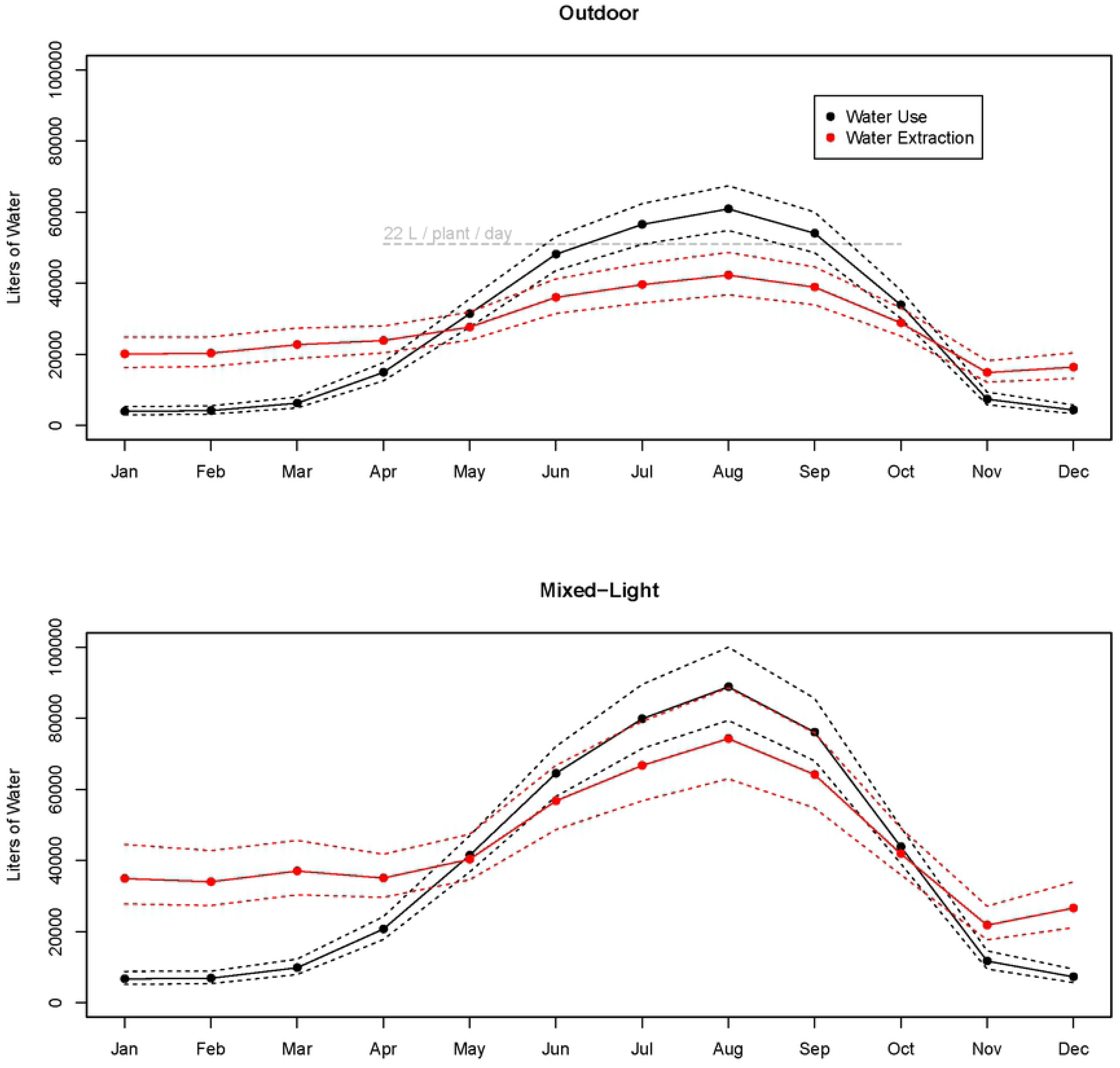
Water Use versus Water Extraction. Predicted monthly water use and water extraction for outdoor and mixed-light operation types. Model estimates are provided for median farm size (cultivation area = 1,098 m^2^). Dashed lines depict 95% confidence intervals for the mean estimate. The rate of 22 L / plant (15 m^2^) / day, which equates to 51,020 L per month for the median farm size of 1,098 m^2^ of cultivation area, is plotted to provide contextual comparison.

Simultaneously estimating the factors influencing *water use* (and *water extraction*) required model fitting to account for zero-inflated, over-dispersed data. Two hurdle models were fit to produce monthly estimates of the response variables *water use* and *water extraction*, respectively, using R statistical programming software [30]. Traditionally used for count data, hurdle models are two-component (binary and continuous) models applied to data with excessive zero values, with the assumption that zero values arise from a process separate from the non-zero values [31–32]. In this context we assume that *water use* is zero only when cannabis cultivation is not occurring and *water extraction* is zero only when cannabis cultivation is not occurring, or when water is being applied only from storage. Non-zero observations are qualitatively distinct from observations of zero, given that if water use or extraction occurs (i.e. non-zero), there is a large minimum amount (∼3,000 L) instead of observations declining linearly to zero. The binary component model of the hurdle model (predicting zero vs non-zero) produces a likelihood of *water use* (and *water extraction*), while the continuous component model produces values for strictly non-zero estimates. The product of the binary and continuous components’ estimates are the full hurdle model predictions for *water use* and *water extraction*, conditional on the likelihood of an observation being non-zero. Using this approach, the hurdle model is able to simultaneously account for monthly observations in which some farms did not use or extract water, while not allowing these observations to artificially reduce the estimates for farms, overall, during said month.

The first (binary) component (referred to hereafter as *binomial model*) of the hurdle models fit a multilevel logistic regression to the binomial response *p_m,t_* indicating whether *water use* or *water extraction* were zero. The predictors included scaled (to standard Z-score) *cultivation area* (*Z*), *month* (*m*), and *operation type* (*t*), and interactions for *cultivation area* and *month*, as well as *cultivation area* and *operation type*. The form of this logistic regression is a generalized linear model (GLM) with a logit link function and *p* being drawn from the binomial distribution:

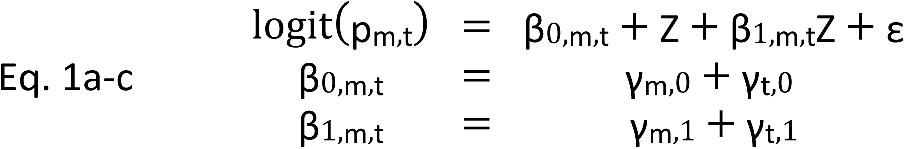

*γ_m,0_* and *γ_t,0_* sum up to the intercept β*_0,m,t_* for a given month and operation type, respectively. *γ_m,1_* and *γ_t,1_* sum up to the slope β*_1,m,t_* for a given month and operation type, respectively. Given twelve months and three operation types, the logistic model contained 36 levelss. With the same predictors as the binomial model, the second (continuous) component model fits a GLM with a Gamma distributed response variable *μ* using an inverse link function:

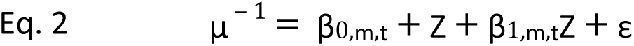

Intercepts and slopes were as described in Eq. 1a-c. The continuous component excluded data with zero *water use* or zero *water extraction*. Hurdle model estimates (*W*) were calculated as the product of the two estimated responses for the binomial model and continuous component model:

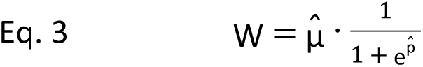

Because model fitting required the use of a link function to account for non-normally distributed data, estimated relationships between continuous variables (i.e. *cultivation area* and *water use*, *cultivation area* and *water extraction*) were non-linear. For the purposes of model interpretation, predictions (liters of *water use* and *water extraction*) were made for each level of the categorical variables (*month* and *operation type*) at the median value of the continuous variable, *cultivation area* (1,098 m^2^). These results can be interpreted as the predicted *water use* or *water extraction* on the median sized farm. Responses in GLMs at the level of the linear predictors are asymptotically normal, and confidence intervals for model responses were calculated with the t-distribution and standard errors. Standard errors for GLMs are the square root of the diagonal elements of the model covariance matrix, estimated via maximum likelihood estimation.

### Storage Capacity Sufficiency

For each farm, the total amount of reported *water use* from April through October was subtracted from the reported storage capacity of the farm to create the response variable *storage balance* (i.e. positive values indicate sufficiency while negative values indicate insufficiency). Extreme outliers of *storage balance* were identified (6 x Interquartile Range) and excluded (n=17). To simultaneously estimate the factors influencing *storage balance* (*S*), a multilevel linear model was fit using the predictors: *cultivation area* (*A*), *operation type* (*t*), *storage type (*g), *source type* (*r*) and an interaction between *cultivation area* and *operation type*:

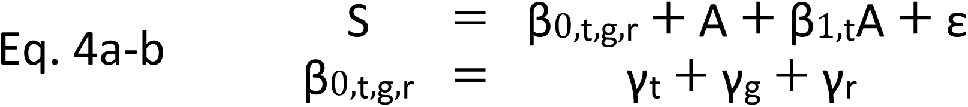

*γ*_t_, *γ*_g_, and *γ*_r_ are components summing to the intercept β*_0,t,g,r_* for a given combination of *operation type*, *storage type*, and *source type*, respectively, resulting in 12 levels total in the model.

### Water Storage, Sources, and Extraction Patterns

An additional hurdle model was fit to determine if predictors of *storage balance* also had reliable effects on seasonal patterns of *water extraction*. The original hurdle model for water extraction was supplemented using two additional predictors of *storage balance* (i.e. *source type* and s*torage type*) along with their interactions with *month*. The intercept defined in Eq. 1b was thus revised accordingly:

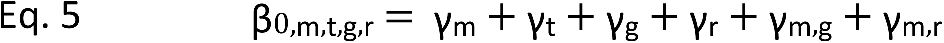

*γ*_m,g_ and *γ*_m,r_ are terms for the interactions of *storage type* and *source type*, respectively, with *month*. The slope term β*_1,m,t_* remained the same. Adding two categorical predictors, with two categories each, resulted in 144 levels in the hurdle models. AIC comparison was used to determine if addition of these parameters was justified [33].

For the purposes of model interpretation, predictions of *water extraction* (liters) were made for each level of the categorical variables (*month*, *operation type*, *source type*, and *storage type*) at the median value of the continuous variable, *cultivation area* (1,098 m^2^).

## Results

The sample of reported data analyzed included outdoor and mixed-light operations, and those with combinations of the two cultivation types (Table 1). Average farm size varied between operation types, with outdoor farms smaller than mixed-light farms and combination farms larger than both. Average reported annual water use and extraction totals were much less for outdoor farms, relative to mixed-light and combination farms; however, there was a notable amount of variation within levels. Average annual water extraction was higher for farms with seasonal water sources than perennial sources, and for farms with ponds relative to those without; although average water storage balances for the forbearance period were greater for these farms with seasonal water sources and ponds.

**Table 1.** Summary Statistics.

|  | Sample size | Mean Cultivation Area (Std Dev) | Mean Annual Water Use (Std Dev) | Mean Annual Water Extraction (Std Dev) | Mean Forbearance Storage Balance (Std Dev) |
| --- | --- | --- | --- | --- | --- |
| **Operation Type** |  |  |  |  |  |
| Outdoor | n = 179 | 1,185 m<sup>2</sup><br>(575) | 358,854 L<br>(303,389) | 371,579 L<br>(322,659) | -72,655 L<br>(279,560) |
| Mixed-light | n = 206 | 1,301 m<sup>2</sup><br>(896) | 533,981 L<br>(458,686) | 602,295 L<br>(508,742) | -38,136 L<br>(522,637) |
| Combination | n = 130 | 1,521 m<sup>2</sup><br>(894) | 500,513 L<br>(344,295) | 531,692 L<br>(370,372) | -54,056 L<br>(428,964) |
| **Source Type** |  |  |  |  |  |
| Seasonal | n = 237 | 1,301 m<sup>2</sup><br>(812) | - | 556,275 L<br>(456,227) | -62,178 L<br>(592,885) |
| Perennial | n = 371 | 1,296 m<sup>2</sup><br>(823) | - | 449,183 L<br>(390,923) | -285,354 L<br>(338,389) |
| **Storage Type** |  |  |  |  |  |
| Pond | n = 61 | 1,438 m<sup>2</sup><br>(807) | - | 831,505 L<br>(534,016) | +384,613 L<br>(591,284) |
| Other | n = 477 | 1,282 m<sup>2</sup><br>(818) | - | 452,254 L<br>(387,702) | -253,927 L<br>(359,426) |

Summary statistics for categorical model parameters. Forbearance storage balance is defined as the reported amount of water used during the pending forbearance period (Apr-Oct) minus the reported storage capacity of the farm.

Models were first developed for two continuous response variables: monthly *water use* and monthly *water extraction*. Model parameters included *cultivation area* as a single continuous predictor variable and two categorical variables: *operation type* and *month*. Model interpretation is reported for both the full hurdle models and the binomial component models. The binomial models estimate the likelihood for *water use* or *water extraction* to occur in a given month, whereas the full hurdle models estimate the amount of monthly *water use* or *water extraction*, conditional on the likelihood of *water use* or *water extraction* occurring for that month (Table 2).

**Table 2.** Binomial Component Models: Water Use and Extraction (Likelihood).

| Month | Outdoor<br>(95% CI) | Mixed-light<br>(95% CI) | Combination<br>(95% CI) |
| --- | --- | --- | --- |
| **Water Use** |  |  |  |
| January | 0.25<br>(0.21, 0.29) | 0.39<br>(0.35, 0.44) | 0.39<br>(0.34, 0.44) |
| February | 0.28<br>(0.24, 0.33) | 0.43<br>(0.39, 0.48) | 0.43<br>(0.38, 0.48) |
| March | 0.37<br>(0.32, 0.41) | 0.53<br>(0.48, 0.58) | 0.53<br>(0.47, 0.58) |
| April | 0.62<br>(0.58, 0.67) | 0.76<br>(0.72, 0.80) | 0.76<br>(0.72, 0.80) |
| May | 0.86<br>(0.82, 0.89) | 0.92<br>(0.90, 0.94) | 0.92<br>(0.90, 0.94) |
| June | 0.99<br>(0.97, 1.00) | 0.99<br>(0.99, 1.00) | 0.99<br>(0.99, 1.00) |
| July | 1.00<br>(0.98, 1.00) | 1.00<br>(0.99, 1.00) | 1.00<br>(0.99, 1.00) |
| August | 1.00<br>(0.98, 1.00) | 1.00<br>(0.99, 1.00) | 1.00<br>(0.99, 1.00) |
| September | 0.98<br>(0.96, 0.99) | 0.99<br>(0.98, 0.99) | 0.99<br>(0.98, 0.99) |
| October | 0.91<br>(0.88, 0.93) | 0.95<br>(0.93, 0.96) | 0.95<br>(0.93, 0.96) |
| November | 0.38<br>(0.33, 0.42) | 0.54<br>(0.49, 0.58) | 0.53<br>(0.48, 0.59) |
| December | 0.27<br>(0.23, 0.31) | 0.42<br>(0.37, 0.47) | 0.42<br>(0.36, 0.47) |
| **Water Extraction** |  |  |  |
| January | 0.45<br>(0.41, 0.50) | 0.52<br>(0.48, 0.57) | 0.53<br>(0.48, 0.58) |
| February | 0.48<br>(0.44, 0.53) | 0.56<br>(0.51, 0.60) | 0.56<br>(0.52, 0.61) |
| March | 0.55<br>(0.51, 0.60) | 0.62<br>(0.58, 0.66) | 0.63<br>(0.58, 0.68) |
| April | 0.69<br>(0.64, 0.73) | 0.75<br>(0.71, 0.78) | 0.75<br>(0.71, 0.79) |
| May | 0.77<br>(0.73, 0.81) | 0.82<br>(0.78, 0.85) | 0.82<br>(0.79, 0.85) |
| June | 0.82<br>(0.78, 0.85) | 0.86<br>(0.83, 0.88) | 0.86<br>(0.83, 0.89) |
| July | 0.80<br>(0.76, 0.84) | 0.84<br>(0.81, 0.87) | 0.85<br>(0.82, 0.88) |
| August | 0.80<br>(0.76, 0.84) | 0.84<br>(0.81, 0.87) | 0.85<br>(0.82, 0.88) |
| September | 0.81<br>(0.78, 0.85) | 0.85<br>(0.82, 0.88) | 0.86<br>(0.83, 0.89) |
| October | 0.80<br>(0.76, 0.83) | 0.84<br>(0.81, 0.87) | 0.84<br>(0.81, 0.87) |
| November | 0.50<br>(0.45, 0.54) | 0.57<br>(0.53, 0.61) | 0.57<br>(0.53, 0.63) |
| December | 0.43<br>(0.39, 0.48) | 0.50<br>(0.46, 0.55) | 0.53<br>(0.46, 0.56) |

Binomial model estimates of likelihood of water use and water extraction for median size cultivation area. Confidence intervals in parentheses.

The binomial models indicated that the likelihood of *water use* was greatest (>0.85) in the growing season (June – October) for all *operation types* (Table 2) and lowest (< 0.40) in the winter months (November – March). However, likelihood estimates were reliably higher for mixed-light and combination cultivation farms than for outdoor farms (November – March), reflecting *water use* extending further into these off-season months. Likelihood of *water extraction* was reliably higher than *water use* from November – March, indicating that water was more likely to be extracted, but not necessarily used, in the offseason. Correspondingly, although the likelihood of *water use* was at certainty (1.00) in the peak growing season (July and August), the likelihood of *water extraction* in these months was reliably lower (<0.85).

Predicted *water use* volumes from the hurdle models indicated strong seasonal patterns, peaking in the late growing season (Fig 2; S1 Table). *Water extraction* volumes were also greatest in the growing season, but showed less seasonal variation than *water use*. For both outdoor and mixed-light cultivation types, *water extraction* was greater than *water use* between November and April, but was less than *water use* from May to October. Overall, *water use* and *water extraction* totals were higher for mixed-light than for outdoor *operation type* farms of the same (median) size, likely resulting from greater density of plants per m^2^ of *cultivation area*.

Models were next developed for *storage balance* to address the necessity of *water extraction* during the growing season (April – October). Reliable predictors of *storage balance* included *cultivation area* (Estimate = −166.09 L; SE = 49.44), *source type* (Seasonal Estimate = 114,945 L; SE = 35,246), and *storage type* (Pond Estimate = 599,763 L; SE = 15,420) (Table 3). The model predicted *storage balance* to be insufficient (−278,879 L) for the median size farm (*cultivation area* = 1,098 m^2^) that relied on perennial sources and used tanks or bladders (“Other”) for storage. Farms of this size relying on seasonal water sources were also predicted to have a negative *storage balance* (−163,930 L). Only farms relying on seasonal water sources that had ponds were predicted to have a positive *storage balance* (435,833 L) at the median size of *cultivation area*. In general, *storage balance* decreased with increasing size of *cultivation area* (Fig 3). Farms without ponds were predicted to have an increasingly negative storage balance, although farms with ponds were predicted to have sufficient *storage balance* for sizes up to nearly one acre of cultivation (3,718 m^2^).

**Fig 3.**
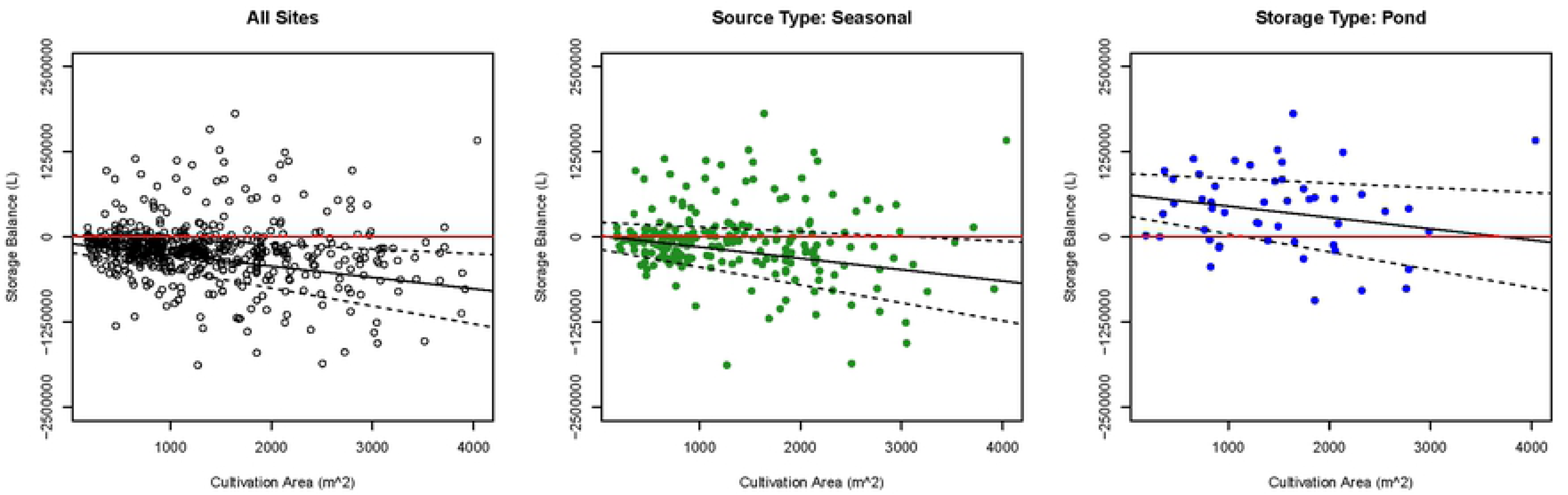
Water Storage Sufficiency. Water storage balance for the cultivation season (April - October) as predicted by cultivation area, source type, and storage type. Reported water use for the cultivation season is subtracted from reported storage capacity, with values of zero indicating storage sufficiency (boundary depicted by red line). Solid lines depict mean estimates, while dashed lines depict 95% confidence intervals.

**Table 3.** Model estimates for water storage balance (Liters).

| Parameter | Estimate | SE |
| --- | --- | --- |
| Intercept | -96,611 | 65,236 * |
| Cultivation Area | -167 | 49 * |
| Operation Type (Mixed-light) | -28,770 | 82,444 |
| Operation Type (Combination) | -160,463 | 92,777 |
| Storage Type Pond | 599,763 | 58,371 * |
| Seasonal Water Source | 114,950 | 35,245 * |
| OT Mixed-Light*Cultivation Area | 28 | 62 |
| OT Combination*Cultivation Area | 131 | 68* |

Estimates for the linear model of water storage balance. Asterisks indicate reliable estimates (95% CI does not overlap zero).

Given the importance of *source type* and *storage type* as predictors of *storage balance*, these parameters were used to refine model estimates of *water extraction*. Both component models of the hurdle model fit with the additional parameters of *source type* and *storage type* were favored by AIC (Binomial AIC = 7,456; Gamma AIC = 119,974) over the original component models (Binomial AIC = 8,265; Gamma AIC = 120,352). The binomial model predicted a reliably higher likelihood of *water extraction* for farms relying on seasonal water sources in the months of January, February, and March relative to farms with at least one perennial water source (Table 4). The pattern was reversed in the summer months of July, August, and September, with farms extracting from seasonal water sources predicted to have a reliably smaller likelihood of *water extraction* than farms with a perennial *source type*. Similarly, the binomial model predicted a reliably higher likelihood of *water extraction* for farms with ponds in the months of January, February, and March, relative to farms without ponds (i.e. *storage type*: other; Table 4). The pattern was reversed in the summer months of July, August, and September, with farms using ponds predicted to have a reliably smaller likelihood of extracting water, relative to farms without ponds.

**Table 4.** Binomial Models: Additional Predictors of Water Extraction (Likelihood).

| Month | Source: Seasonal<br>Storage: Other<br>(95% CI) | Source: Perennial<br>Storage: Other<br>(95% CI) | Source: Seasonal<br>Storage: Pond<br>(95% CI) |
| --- | --- | --- | --- |
| January | 0.59<br>(0.51, 0.66) | 0.31<br>(0.26, 0.36) | 0.84<br>(0.71, 0.91) |
| February | 0.62<br>(0.54, 0.69) | 0.34<br>(0.30, 0.40) | 0.82<br>(0.69, 0.90) |
| March | 0.65<br>(0.57, 0.72) | 0.44<br>(0.38, 0.49) | 0.84<br>(0.72, 0.92) |
| April | 0.72<br>(0.64, 0.78) | 0.63<br>(0.57, 0.68) | 0.83<br>(0.70, 0.92) |
| May | 0.71<br>(0.63, 0.78) | 0.81<br>(0.76, 0.85) | 0.53<br>(0.40, 0.66) |
| June | 0.64<br>(0.56, 0.72) | 0.96<br>(0.93, 0.97) | 0.34<br>(0.23, 0.48) |
| July | 0.60<br>(0.52, 0.68) | 0.95<br>(0.92, 0.97) | 0.36<br>(0.25, 0.50) |
| August | 0.61<br>(0.53, 0.68) | 0.95<br>(0.92, 0.97) | 0.31<br>(0.20, 0.44) |
| September | 0.62<br>(0.54, 0.70) | 0.96<br>(0.93, 0.98) | 0.34<br>(0.23, 0.48) |
| October | 0.65<br>(0.57, 0.72) | 0.89<br>(0.85, 0.92) | 0.53<br>(0.39, 0.67) |
| November | 0.50<br>(0.42, 0.57) | 0.45<br>(0.40, 0.51) | 0.60<br>(0.47, 0.72) |
| December | 0.50<br>(0.42, 0.58) | 0.32<br>(0.27, 0.38) | 0.77<br>(0.63, 0.86) |

Water extraction model estimates for median size cultivation area, with additional predictors of source type and storage type. Confidence intervals in parentheses.

The volume of *water extraction* predicted by the full hurdle model followed the pattern of the binomial model (Fig 4; S2 Table). *Water extraction* totals were reliably greater for farms with seasonal water sources in the months of January (0.59), February (0.62), and March (0.65) relative to farms with at least one perennial water source (0.31, 0.34, and 0.44, respectively; Table 4). The pattern was reversed in the summer months of June, July, August, and September, with predicted amount of *water extraction* from farms with a seasonal *source type* lower than farms with a perennial *source type*. Farms with ponds demonstrated an even more pronounced divergence from farms using perennial sources. *Water extraction* totals were reliably higher for farms with pond *storage type* in the months of January (0.84), February (0.82), and March (0.83) relative to other (tanks or water bladders) *storage type*, regardless of *source type*. The pattern was reversed in the summer months of July, August, and September, with predicted amount of *water extraction* from farms with ponds lower than from farms without ponds, regardless of *source type*. However, this difference was only reliable between farms with ponds and those without, in which the *source type* was perennial, as 95% confidence intervals overlapped when comparing farms with and without ponds, in which *source type* was seasonal.

**Fig 4.**
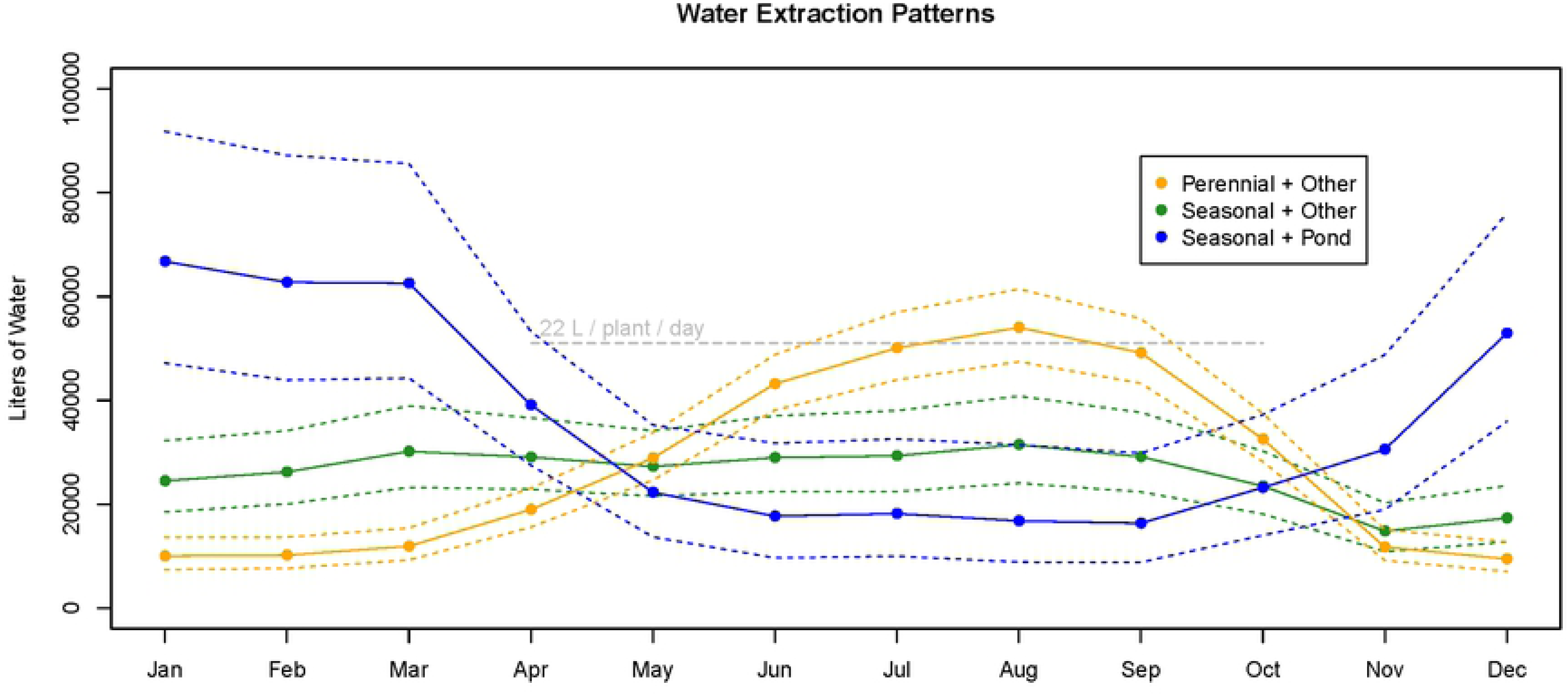
Additional Predictors of Water Extraction. Monthly water extraction, based on source type and storage type. Model estimates are provided for median farm size (cultivation area = 1,098 m^2^). Dashed lines depict 95% confidence intervals for the mean estimate. The rate of 22 L / plant (15 m^2^) / day, which equates to 51,020 L per month for the median farm size of 1,098 m^2^ of cultivation area, is plotted to provide contextual comparison.

## Discussion

Cannabis cultivation has been considered an emerging environmental threat to Northern California watersheds [5]. While there is strong evidence that a large number of farms are located in sensitive and remote locations [4], until now, there had been little data about their actual water demand patterns. Applying newly available data, we modeled the characteristics of water extraction, storage, and use for over 600 cannabis farms in Northern California, providing policy relevant information on these patterns.

We found reliable variation between months in terms of both water use and water extraction. For all operation types, water extraction in offseason months exceeded water use, reflecting input to storage rather than immediate use for cultivation. This stored water likely reduced the need to withdraw water in summer months, as water extraction was less than water use during this period. However, farms did not generally have enough storage to completely refrain from extracting from April through October. The same useful predictors of storage sufficiency (type of storage infrastructure and seasonality of water sources) had reliable effects on extraction patterns, further emphasizing that patterns of input to storage are linked to storage capacity and whether a farm needs to store water. Farms relying on seasonal water sources, and especially those with ponds, weighted their annual extraction profile toward offseason months, whereas farms incorporating perennial sources had extraction profiles that more closely followed plant demand over the growing season. The results observed herein demonstrate that estimating the water demands of cannabis cultivation will require accounting for monthly extraction patterns, in addition to cultivation strategies and farm characteristics that influence them. Furthermore, given the link between water storage and extraction patterns, widespread storage insufficiency represents an important topic of discussion in light of future natural (e.g. drought) and regulatory (e.g. forbearance) restrictions on seasonal water sources.

### Storage Insufficiency

The results suggest that many farms may need to expand water storage capacity if they are to eliminate the need for surface water extractions during the growing season. Beginning in 2019, forbearance requirements will be implemented by the California State Water Resources Control Board that prohibit extraction from surface water (and springs that deliver to surface water) from April through October. Therefore, although farms included in the current study were not subject to these restrictions at the time data were collected, farms relying on surface water (and connected springs) will be required to either develop storage or seek an alternative water source, such as subsurface water. Furthermore, the data analyzed in the current study were collected after a particularly wet winter (2016-2017) [34] and many seasonal water sources reported, herein, may not be available during drought, or even normal years. While farms may have the options of developing storage for surface water and/or rain catchment, receiving water from offsite, or extracting subsurface water, previous work has suggested that drilling wells may be the method of choice to source water in a manner that will provide insurance against drought and comply with forbearance requirements [10]. The appeal of drilling a well may reflect difficulties associated with obtaining storage infrastructure, which could be partially responsible for this decision.

Although farms with ponds generally had sufficient water storage to comply with forbearance requirements, only approximately 10% of farms reported use of a pond for cannabis irrigation. There are logistical, financial, environmental, and regulatory concerns that are likely limiting this option for farms. Aside from the costs and engineering constraints for building ponds on rugged terrain, there may be difficulty in ensuring ponds are not situated on seasonal watercourses, thus capturing streamflow and rendering them non-compliant with state and county regulations. Depending on where they are located, ponds may also serve as habitat for invasive species, such as bullfrogs, which are also of concern to regulatory agencies. Although water storage tanks could avoid these concerns, the costs of units themselves and the availability of appropriate terrain to site numerous large water tanks may pose complications for farms in rugged terrain. With increasingly larger farms in such areas, the likelihood of securing enough tanks to meet water needs becomes increasingly smaller. Under these circumstances, not all farms that rely on seasonal water may be able to meet forbearance requirements (or outlast drought conditions), due to a lack of water storage. In these cases, farmers may instead choose to bypass storage requirements by drilling wells, which emphasizes the need to account for extraction patterns of perennial versus seasonal water sources.

### Water Sources and Ecological Impacts

Based on results observed in the current study, farms using wells would be expected to follow an extraction pattern that matches plant demand, overlapping with diminishing instream flow during summer dry months [35]. It is known that extraction of ground water may have a delayed impact on instream flow on the order of weeks, months, or years, depending on the depth of extraction, conductivity of the soil, and the recharge received from precipitation [36]. As a result, understanding lagged effects on instream flow will be useful when assessing the potential benefits of shifting the instream flow impacts of cannabis water extraction out of the crucial summer drought months. An accurate assessment of the benefits and risks of well extraction will require a better understanding of the geology and hydrology in areas where cannabis cultivation occurs and on the spatial and temporal dimensions of groundwater-surface water interactions [37–38]. While there may be benefits of lagged impacts of wells on instream flow, the possibility of wells instead being directly hydrologically connected to streams may result in additional concerns for instream flow [39].

Wells that are shallow and close to surface water have a high likelihood of directly capturing stream flow [40–41]. As a result, water extraction would have a minimal lag on instream impacts and the extraction pattern, matching plant demand, would directly overlap with the most crucial low instream flow period. Further work is needed to determine the propensity for wells servicing cannabis farms to be located near streams and the degree to which they are hydrologically connected. For wells that are determined to be capturing surface water, forbearance requirements will prohibit the use of these sources from April through October. The ability of these farms to switch to storing water or to drill a new well would then influence their ability to remain in compliance with regulations. For sites that are currently outside of the regulated industry, this may be a barrier to becoming permitted. Given the link between water sources and seasonal extraction patterns demonstrated in the current study, it will be useful to determine how unpermitted sites (i.e. those operating outside the regulated industry) may use water in order to develop a holistic understanding of the impact of cannabis cultivation in general on instream flow.

Although the current study demonstrated that summer water extraction is reduced for farms that use seasonal water sources, unpermitted sites frequently use seasonal sources opportunistically during the summer growing season [18]. In fact, illegal diversions are a major issue, given that the majority of cannabis cultivation in the North Coast of California is currently unpermitted [42]. In those cases, plant demand (i.e. *water use*) estimates provided herein may be more appropriate predictors of water impacts, assuming little to no storage is being used. However, it is difficult to anticipate what proportion of these farmers incorporate water storage, either due to necessity or concern for environmental impacts. This simultaneously emphasizes the importance of these sites entering the regulated industry [43] and illustrates the limitations of trying to estimate collective impacts of cannabis cultivation without sufficient data on cultivation practices of unpermitted operations.

### Future Research Needs

A lack of understanding of illicit (i.e., unpermitted) cannabis farming practices represents one of several limitations of this study and a need for additional data. Field observations from warrant inspections on unpermitted cannabis farms have revealed several cultivation practices that may affect how water is extracted, stored, and used for cannabis. For example, perennial springs that would otherwise feed small streams are often dammed by cannabis cultivators to store water for critical summer months. Alternatively, spring diversions often feed directly into storage tanks without overflow protections, thereby moving water out of its regular channel and dispersing it in upland areas. Empirical streamflow studies may be useful to assess the impact of these practices, comparing expected water extraction totals to instream flow reductions in a paired watershed design. These efforts would be aided by improving water extraction estimates themselves, using more detailed data to improve accuracy.

Data collection incorporating additional parameters that influence water use for cannabis cultivation would be beneficial to both regulators and farmers. The results of this study indicate significant differences in predicted water use and extraction amounts as a result of operation types known to differ in plant sizes, spatial arrangement, and evapotranspiration potential based on ambient temperature and humidity. However, the precise relationship between these variables remains unknown. Furthermore, there are certainly additional factors, such as the soil type, local climate, and cultivar that will influence water consumption [26]. A better understanding of these factors could potentially inform water conservation best practices targeted toward specific cultivation strategies and growing conditions, the variety of which are a hallmark of the cannabis industry in Northern California. Improved estimates that account for diverse cultivation practices may also help growers to know how their use compares with the expected range of water use and thus be able to identify and address operational inefficiencies.

## Conclusion

This study demonstrates that predicting water demands of cannabis farms requires consideration of the seasonal patterns of water extraction, cultivation practices, water sources, and storage availability. Pending decisions for farmers aiming to comply with regulations may influence these seasonal extraction patterns and in turn, inform relative impacts to instream flow. In general, more data are needed on cultivation practices to help determine additional factors that influence water demand by cannabis farms. Regulators and researchers may continue to explore the geographical, climatic, and operation-specific factors that influence water demand and more specifically tailor regulations based on these factors. Cannabis farmers may benefit from an established understanding of what water use expectations are and should be. All stakeholders will benefit from determinations of environmental impacts, so that regulatory objectives can be effective, transparent, and achievable [44–45].

## Acknowledgements

We are grateful for feedback from D. Kuszmar, B. McFadin, and J. Carah, whose review and comments improved the quality of the manuscript. We thank J. Stapp for his help with data acquisition. We also thank NCRWQCB staff responsible for implementation of the North Coast Regional Water Quality Control Board Cannabis Waste Discharge Regulatory Program, which made this work possible.

## Supporting Information

**S1 Figure.**
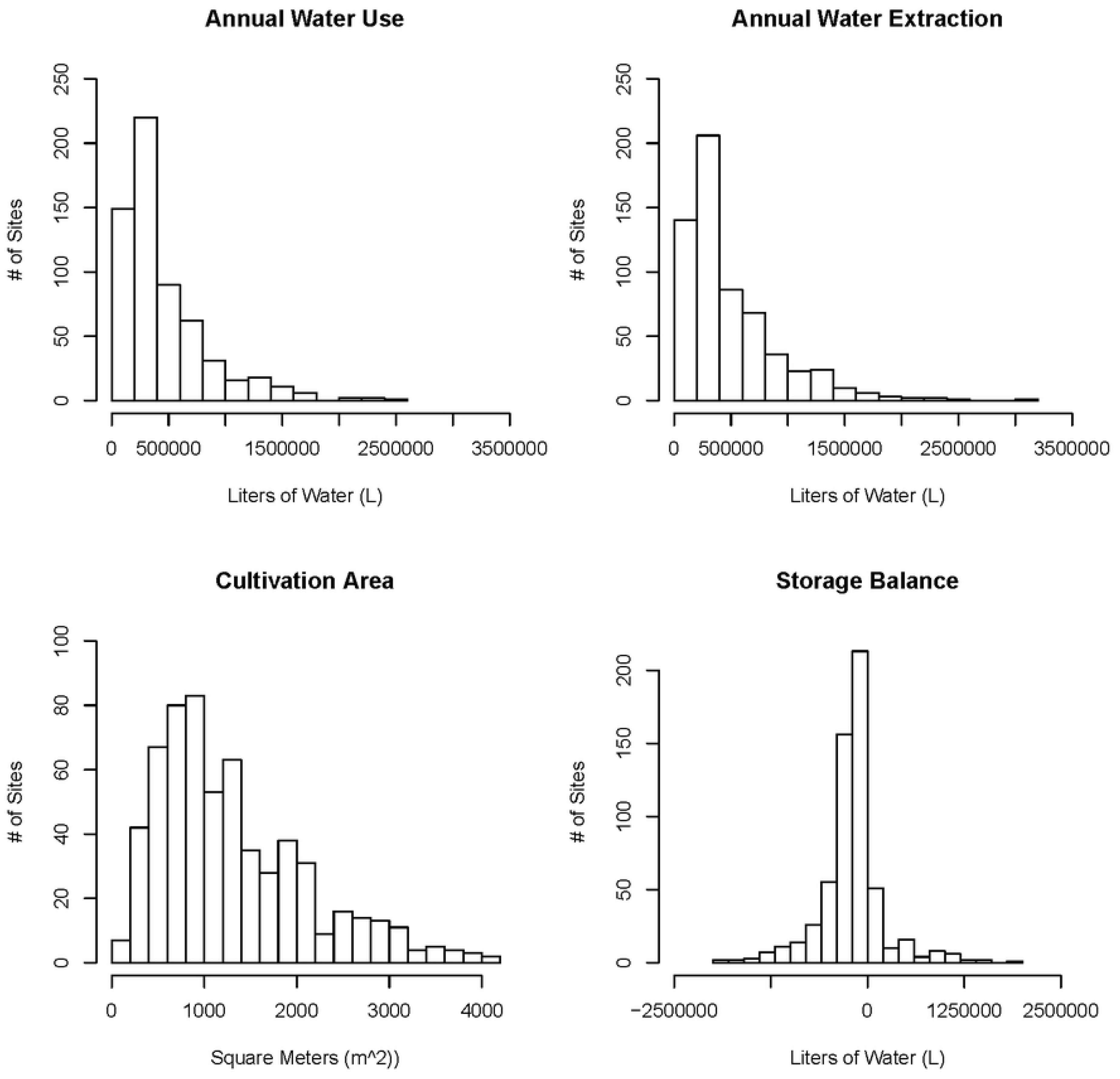
Distributions of Summary Statistics. Summary statistics for the continuous model parameters of *cultivation area* (predictor) and *storage balance* (response). Annual water use and annual water extraction are depicted for descriptive purposes only and are not included as model predictors or response variables.

**S2 Figure.**
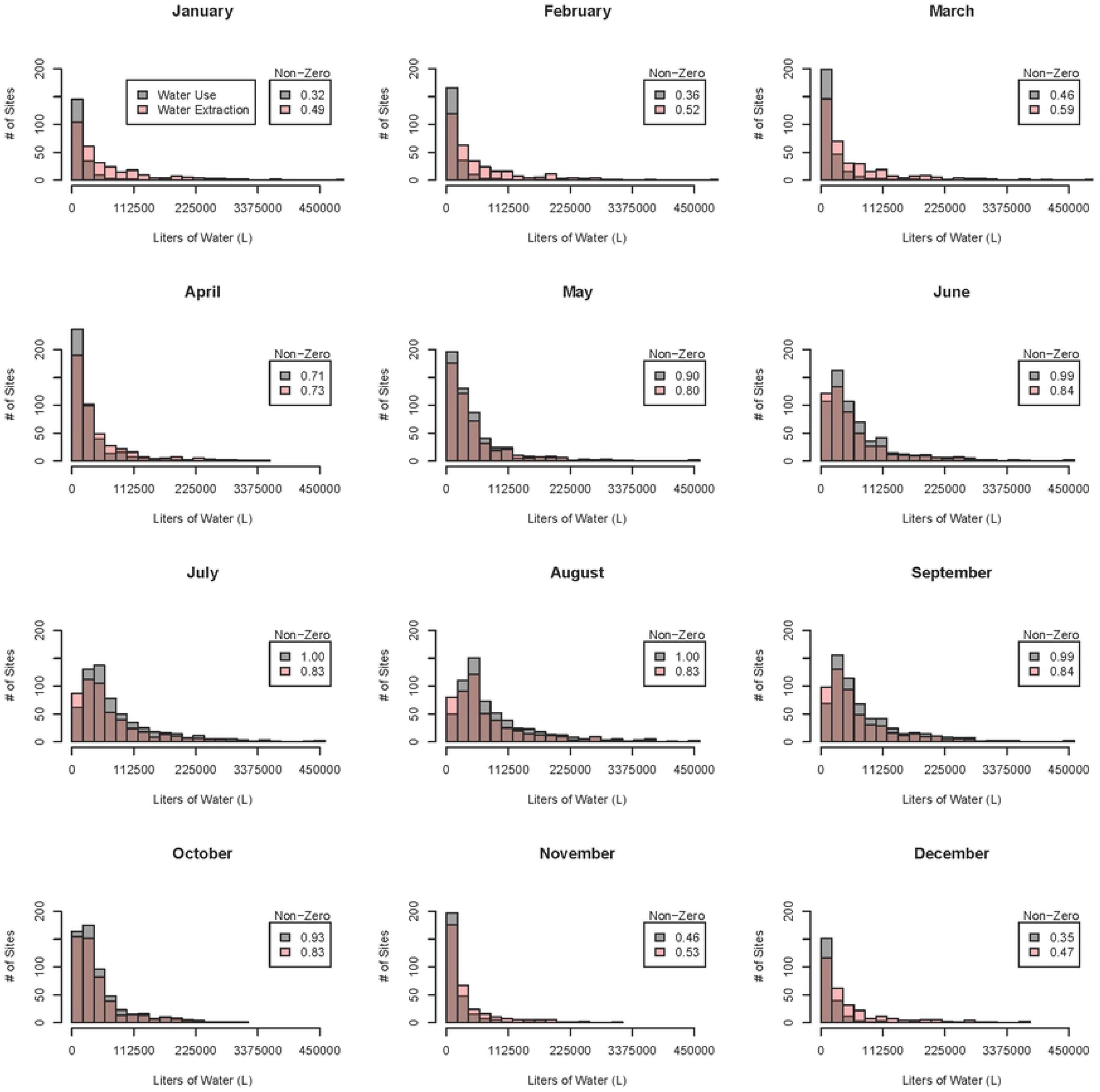
Monthly Water Data Distributions. Raw monthly water use and water extraction values. Distributions depict non-zero observations, used in the continuous (gamma) model component of the hurdle model. The proportion of monthly observations that were non-zeros is also provided, corresponding to binary input to the binomial model component of the hurdle model.

**S1 Table.** Full Hurdle Model Estimates (Liters).

| Month | Outdoor<br>(95% CI) | Mixed-light<br>(95% CI) | Combination<br>(95% CI) |
| --- | --- | --- | --- |
| **Water Use** |  |  |  |
| January | 3964<br>(2984, 5320) | 6720<br>(5180, 8827) | 6664<br>(5061, 8857) |
| February | 4193<br>(3207, 5529) | 6897<br>(5407, 8887) | 6843<br>(5290, 8917) |
| March | 6278<br>(4950, 8000) | 9857<br>(7965, 12288) | 9746<br>(7780, 12261) |
| April | 14926<br>(12573, 17705) | 20729<br>(17792, 24235) | 20284<br>(17302, 23815) |
| May | 31419<br>(27656, 35622) | 41482<br>(36788, 6957) | 39882<br>(35221, 45328) |
| June | 48163<br>(43557, 53090) | 64492<br>(57992, 72061) | 60895<br>(54388, 68615) |
| July | 56529<br>(50950, 62368) | 79827<br>(71485, 89513) | 74403<br>(65954, 84505) |
| August | 60890<br>(54845, 67400) | 88827<br>(79399, 99965) | 82161<br>(72532, 93934) |
| September | 54050<br>(48590, 60022) | 76064<br>(68021, 85574) | 71090<br>(62988, 80903) |
| October | 33900<br>(30139, 38048) | 43854<br>(39125, 49361) | 42107<br>(37428, 47578) |
| November | 7404<br>(5859, 9395) | 11738<br>(9504, 14602) | 11571<br>(9258, 14523) |
| December | 4369<br>(3325, 5794) | 7305<br>(5693, 9476) | 7239<br>(95561, 9498) |
| **Water Extraction** |  |  |  |
| January | 20092<br>(16302, 24861) | 34932<br>(27787, 44527) | 31408<br>(25028, 39786) |
| February | 20314<br>(16622, 24908) | 34010<br>(27366, 42778) | 30850<br>(24860, 38587) |
| March | 22727<br>(18892, 27399) | 37038<br>(30346, 45657) | 33501<br>(27495, 41083) |
| April | 23882<br>(20410, 27963) | 35070<br>(29649, 41771) | 32215<br>(27320, 38159) |
| May | 27660<br>(23999, 31881) | 40341<br>(34560, 47397) | 36611<br>(31495, 42755) |
| June | 36022<br>(31494, 41204) | 56753<br>(48657, 66744) | 49482<br>(42612, 7840) |
| July | 39602<br>(34473, 45513) | 66701<br>(56779, 79112) | 56693<br>(48424, 66911) |
| August | 42264<br>(36735, 48660) | 74263<br>(62977, 88517) | 61973<br>(52726, 73514) |
| September | 38898<br>(33941, 44596) | 64136<br>(54737, 75862) | 54901<br>(47041, 64578) |
| October | 28833<br>(25143, 33064) | 41876<br>(35976, 49080) | 37881<br>(32691, 44121) |
| November | 14877<br>(12204, 18194) | 21815<br>(17699, 27162) | 20787<br>(16885, 25755) |
| December | 16404<br>(13240, 20413) | 26623<br>(21133, 33988) | 24716<br>(19656, 31349) |

Water use and extraction model estimates for median size cultivation area, by operation type. Confidence intervals in parentheses.

**S2 Table.** Full Hurdle Model Estimates for Additional Predictors (Liters).

| Month | Source: Perennial<br>Storage: Other<br>(95% CI) | Source: Seasonal<br>Storage: Other<br>(95% CI) | Source: Seasonal<br>Storage: Pond<br>(95% CI) |
| --- | --- | --- | --- |
| January | 9991<br>(7386, 13640) | 24482<br>(18514, 32237) | 66754<br>(47181, 91822) |
| February | 10158<br>(7628, 13630) | 26205<br>(20010, 34149) | 62758<br>(43894, 87217) |
| March | 11896<br>(9226, 15410) | 30162<br>(23227, 38946) | 62561<br>(44305, 85627) |
| April | 18956<br>(15565, 23061) | 29052<br>(22858, 36603) | 39124<br>(27504, 53266) |
| May | 28877<br>(24634, 33778) | 27256<br>(21542, 34154) | 22268<br>(13672, 35255) |
| June | 43182<br>(38113, 48807) | 28963<br>(22447, 37013) | 17684<br>(9649, 31735) |
| July | 50087<br>(43948, 56981) | 29316<br>(22418, 38029) | 18167<br>(9971, 32552) |
| August | 54027<br>(47448, 61430) | 31459<br>(24057, 40821) | 16770<br>(8851, 31444) |
| September | 49173<br>(43291, 55747) | 29136<br>(22355, 37667) | 16331<br>(8804, 29828) |
| October | 32524<br>(28173, 37455) | 23488<br>(18112, 30224) | 23193<br>(14003, 37241) |
| November | 11730<br>(9154, 15092) | 14812<br>(10840, 20250) | 30554<br>(18948, 48788) |
| December | 9431<br>(7035, 12745) | 17312<br>(12715, 23548) | 52948<br>(35952, 76012) |

Water extraction model estimates for median size cultivation area, with additional predictors of source type and storage type.
Confidence intervals in parentheses.

